# Distinguishing the neural correlates of perceptual awareness and post-perceptual processing

**DOI:** 10.1101/2020.01.15.908400

**Authors:** Michael A. Cohen, Kevin Ortego, Andrew Kyroudis, Michael Pitts

## Abstract

To identify the neural correlates of perceptual awareness, researchers often compare the differences in neural activation between conditions in which an observer is or is not aware of a stimulus. While intuitive, this approach often contains a critical limitation: In order to link brain activity with perceptual awareness, observers traditionally report the contents of their perceptual experience. However, relying on observers’ reports is problematic because it is difficult to know if the neural responses being measured are associated with conscious perception or with post-perceptual processes involved in the reporting task (i.e., working memory, decision-making, etc.). To address this issue, we combined a standard visual masking paradigm with a recently developed “no-report” paradigm in male/female human participants. In the visual masking paradigm, observers saw images of animals and objects that were visible or invisible depending on their proximity to masks. Meanwhile, on half of the trials, observers reported the contents of their perceptual experience (i.e., report condition), while on the other half of trials they refrained from reporting about their experiences (i.e., no-report condition). We used electroencephalography (EEG) to examine how visibility interacts with reporting by measuring the P3b event related potential (ERP), one of the proposed canonical “signatures” of conscious processing. Overall, we found a robust P3b in the report condition, but no P3b whatsoever in the no-report condition. This finding suggests that the P3b itself is not a neural signature of conscious processing and highlights the importance of carefully distinguishing the neural correlates of perceptual awareness from post-perceptual processing.

**Significance statement:** What are the neural signatures that differentiate conscious and unconscious processing in the brain? Perhaps the most well-established candidate signature is the P3b event-related potential (ERP), a late slow wave that appears when observers are aware of a stimulus, but disappears when a stimulus fails to reach awareness. Here, however, we found that the P3b does not track what observers are *perceiving* but instead tracks what observers are *reporting*. When observers are aware of simple visual stimuli, the P3b is nowhere to be found unless observers are reporting the contents of their experience. These results challenge the well-established notion of the P3b as a neural marker of awareness and highlight the need for new approaches to the neuroscience of consciousness.

## Introduction

A long-standing debate in cognitive neuroscience focuses on the neural correlates of perceptual awareness. In recent years, two dominant classes of theories have emerged. One group of theories believe information reaches awareness when it is globally accessible to various systems including memory, language, and action-planning (Dehaene & Naccache, 2001; Baars, 2002; Cohen & Dennett, 2011; Cohen et al., 2012), or when it is indexed by higher-order, reality-monitoring mechanisms (Lau, 2019; Brown et al., 2019). Under these views, subconscious processing occurs in sensory regions, such as the ventral visual pathway (Dehaene et al., 2001; Sergent et al., 2005;), while conscious processing occurs within domain-general fronto-parietal networks (Odegaard et al., 2017; Dehaene, et al., 2017).

Meanwhile, another group of theories believe the correlates of consciousness are found within sensory regions of the brain. For example, in the case of vision, these theories maintain that conscious processing occurs (Lamme et al., 2003; 2010), or perceptual experience is instantiated by the grid-like structure (Koch et al., 2016; Tononi et al., 2016; Boly et al., 2017), within posterior cortical and thalamo-cortical feedback loops between regions like LGN, V1, V4, and IT cortex. According to these views, recurrent processing, or cause-effect power within posterior cortex, causes information to reach awareness even if that information is never accessed by higher-order cognitive mechanisms.

How do sensory theories account for the fact that visible stimuli reliably activate frontoparietal regions while invisible stimuli only activate sensory regions? Generally speaking, sensory theorists believe these results conflate the neural responses associated with perceptual awareness with the neural responses associated with post-perceptual processing (Aru et al., 2012). In nearly all prior studies, observers were tasked with making judgments about stimuli that they did or did not consciously perceive (e.g., Was the target an animal or object?). However, when stimuli are not perceived, such judgments cannot be made and observers can only report that they did not see anything or provide a random guess. Therefore, the “aware” condition includes additional steps in post-perceptual processing that the “unaware” condition does not (e.g., categorization, short-term memory, decision-making, etc.). For this reason, researchers recently developed “no-report” paradigms in which observers do not make any judgments about their perceptual experience (Frässle et al., 2014; Shafto & Pitts, 2015; Tsuchiya et al., 2015; Koch et al., 2016; Wiegand et al., 2018; Farooqui & Manley, 2018; Kapoor et al., 2020).

To date, however, studies incorporating no-report conditions have used non-standard manipulations of awareness and/or passive viewing conditions, which have tempered enthusiasm and confidence in their results. Thus, in this study we combined a classic visual masking paradigm with a novel no-report paradigm. Specifically, we compared neural responses to visible versus invisible stimuli when observers did and did not report their experiences. We focused on arguably the largest and most replicable piece of neural evidence cited by cognitive theories: the P3b event-related potential (ERP). While numerous signatures of conscious processing have been proposed (i.e., an “ignition” of fronto-parietal circuits, an increase in long-range neural synchrony, and a late burst of gamma-band oscillations, Dehaene, 2014), the P3b has been widely replicated and continues to be routinely cited as a neural marker of conscious experience (Railo et al., 2011; Naccache, 2018; Ye et al., 2019; Derda et al., 2019).

Overall, we found that when observers reported their experiences, a large P3b separated visible and invisible stimuli. When observers were shown the exact same stimuli, but did not make any reports, the P3b disappeared completely. These results provide the most compelling evidence to date that the P3b is not a signature of conscious processing and is instead linked with post-perceptual processing. It should be stressed, however, that while our results conclusively rule-out one of the main proposed signatures of conscious perception, they do not in any way disprove cognitive theories of consciousness. Indeed, those theories may ultimately be correct even if the P3b is not a true signature of conscious awareness.

## Materials and Methods: Experiment 1

The masking paradigm was closely modeled on the seminal experiments of Dehaene et al. (2001). Our experimental design and planned analyses were pre-registered prior to data collection on the Open Science Framework: https://osf.io/rgfxy/.

### Participants

Thirty three participants from the Reed College community completed the experiment in order to obtain the pre-registered sample size of twenty participants after exclusions (see exclusions section below). All participants had normal or corrected-to-normal visual acuity, with no head trauma or concussions within the past year and no known history of seizures or epilepsy. Participants were 18-26 years old (mean = 20.7 years), 15 male, 18 female, 32 right-handed. Informed consent was obtained prior to each experimental session and all procedures were approved by the IRB at Reed College (IRB no. 2018-S14).

### Apparatus

Brain electrical activity was recorded using a custom 64-channel electrode cap with equidistantly spaced electrodes (EASYCAP). Electrode positions reported here refer to the nearest channels of the international 10-20 system. Electrode impedances were kept <10kΩ, and signals were amplified by two 32-channel amplifiers (Brain Amp Standard; Brain Products), band-pass filtered from 0.1 to 150 Hz, and digitized at 500 Hz. Eye movements and blinks were monitored by left and right horizontal EOG channels and a vertical EOG channel under the left eye. An electrode attached to the right mastoid served as the reference during recording.

### Stimuli

The critical stimuli were 32 line drawings, 16 objects and 16 animals. Each participant was shown all 32 stimuli over the course of the experiment. The stimuli were divided into 4 groups (A, B, C, and D), each group consisting of 4 animals and 4 objects. Each participant, according to their subject number, was assigned a stimulus group (A, B, C, or D) for each combination of trial type and task condition: (1) visible, report; (2) masked, report; (3) visible, no-report; and (4) masked, no-report. These stimulus groups were counterbalanced across participants using a Williams Latin square design (Williams, 1949). Therefore, in each task condition (report, no-report), exactly half of the stimuli were presented to each participant. Due to the manipulation of awareness via masking, exactly half of the presented stimuli were visible within each task condition.

The masks were constructed using line segments from the animal and object stimuli, overlaid upon each other such that no obvious shapes could be perceived. A total of 8 mask variants were used (created by rotating and flipping the original mask), and a pair of non-matching masks was randomly selected for each trial. All stimuli and masks were 625×625 pixel (px) images. Large green circles (RGB: 200,255,200) with a diameter of 750 px, served as the target stimuli in the no-report condition.

Stimuli were controlled using Psychophysics Toolbox Version 3 for MATLAB (Kleiner et al., 2007). All stimuli were presented on a white background (BenQ 120Hz monitor, 1920×1080 px). Participants were seated 70 cm from the monitor, thus making the average size of the critical stimuli 8.17°, the masks 14.05°, and the green circles 16.26°. All stimuli were presented at the center of the screen, while participants maintained fixation on a small red fixation dot (0.20°) that was continuously visible throughout the procedure. All of the MATLAB code and image files needed to run the experiment are available on the Open Science Framework: https://osf.io/rgfxy/. In addition, videos of the stimulus presentation sequences are also available on the OSF website and on the first and last authors’ lab websites.

### Experimental Design

Each participant completed all 4 parts of the experimental procedure in the same order: (1) 15 no-report blocks with 64 trials in each block, (2) an incidental memory test on those task-irrelevant stimuli, (3) 15 report blocks with 64 trials in each block, and (4) an incidental memory test on those task-relevant stimuli.

Each trial consisted of a series of two masks, two blanks, and either a critical stimulus or a complementary blank, lasting a total of 633 ms (see Figure 1). Masks were presented for 100 ms, blanks for 200 ms, and the critical stimuli/blanks for 33 ms (closely following Dehaene et al. 2001). On visible trials, the 200 ms blank gaps were presented immediately before and after the presentation of the critical images (i.e., the animal and object line drawings). On the invisible trials, the masks themselves were presented immediately before and after the presentation of the critical images rendering them invisible to observers. In both visible and invisible trials, there was a 1000 ms blank inter-trial interval that occurred after each trial and before the next. On the subset of trials in which a green circle stimulus appeared, its onset within the trial sequence was chosen randomly from 0-1267 ms, and its duration was always 300 ms.

**Figure 1.**
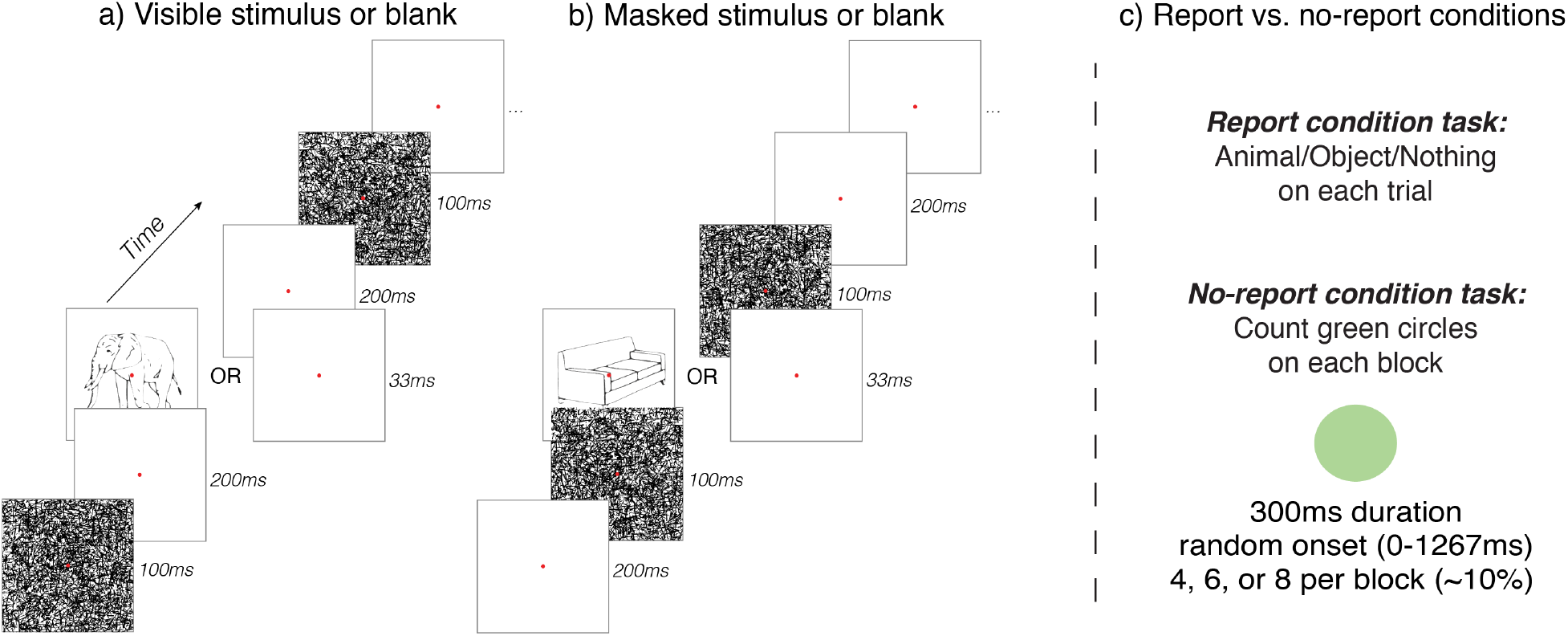
Design of Experiment 1. Stimuli (i.e., animals or objects) or blank displays were presented in between masks. a) On visible trials, there were 200 ms gaps separating the stimuli from the masks. b) On masked trials, the masks came immediately before and after the stimulus, rendering them completely invisible. c) In the report condition, participants reported on a trial-by-trial basis whether they saw an animal, object, or nothing. In the no-report condition, the stimulus presentation sequence was the same, but instead of reporting on these stimuli, participants counted the number of times they saw a green circle and reported their count at the end of each block.

Line drawings of animals/objects were presented on 50% of all trials, with 25% being “visible” (lightly masked) and 25% being heavily “masked” (invisible). Complementary blanks were presented on 50% of all trials, with 25% lightly masked and 25% heavily masked. This was to ensure equal numbers of stimulus and blank trials within each masking condition, for purposes of subtracting mask-blank-mask ERPs from maskstimulus-mask ERPs. Green circles appeared on ~10% of trials on average (4, 6, or 8 in each block). All stimulus parameters were identical across conditions, with the key change between conditions being the task the participants were instructed to complete.

During the no-report condition, participants were instructed to count how many times the green circle appeared during each 64-trial block, providing a 3-alternative-forced-choice (3-AFC) button-press response at the end of each block to indicate 4, 6, or 8 green circles (number of circles presented varied randomly across blocks, and all trials in which a green circle appeared were excluded from ERP analyses). During the report condition, participants were instructed to provide a 3-AFC button-press response after every trial, indicating whether they saw an “animal”, “object”, or “nothing” between the pair of masks.

After the 15 blocks of each condition, participants were given an incidental memory test, to verify the manipulation of visibility via masking, regardless of the task. Accordingly, during these memory tests, participants were shown the 8 “visible” (lightly masked) images, the 8 heavily “masked” (invisible) images, and 8 foil images (never presented), in a random order. While viewing each image (duration unrestricted), participants provided either a “yes” or “no” button-press to indicate whether they remembered seeing that particular line drawing during the preceding blocks of trials.

### Statistical Analysis

EEG data were processed using Brain Vision Analyzer 2.2 (Brain Products). Recordings were re-referenced to the average of the left and right mastoids and low-pass filtered at 25 Hz with a 24 dB/Oct roll-off. The left and right horizontal EOG channels and the vertical EOG channel and electrode FP1 were re-referenced as a bipolar pairs for artifact rejection. Trials containing artifacts (e.g., blinks, eye movements, muscle noise) were rejected semi-automatically using peak-to-peak amplitude thresholds with starting values of 50μV for eye movements, 100μV for blinks, and 150μV for other artifacts, adjusted on a subject-by-subject basis. On average, 17.4% of trials were excluded due to artifacts among included participants, leaving on average ~180 trials for analysis per condition. More specifically, in the report condition, 9.9% of trials were rejected for eye blinks and 1.4% were removed for eye movements. In the no-report condition, 11.8% of trials were rejected for eye blinks and 1.6% of trials were removed for eye movements. The percentage of trials removed for eye blinks and eye movements were not significantly different between the two conditions (eye blinks: *t*(19)=1.272, *p*=0.22, eye movements: *t*(19)=0.813, *p*=0.43).

ERPs were time-locked to the onset of stimuli and blanks, and baseline corrected from - 200 to 0 ms. To isolate ERPs elicited by stimuli from overlapping activity caused by the masks, we used the same mask subtraction procedure of Dehaene et al. (2001). To accomplish this, ERPs elicited by lightly masked and heavily-masked blank trials were subtracted from the ERPs elicited by lightly masked and heavily-masked stimulus present trials, respectively. Time windows and electrodes for statistical analysis were preregistered and based on typical spatial-temporal regions of interest (ROIs) for the P3b, which is maximal over centro-parietal electrode sites between 300-600 ms (Sergent et al., 2005; Azizian et al., 2006; Del Cul et al., 2007; Polich, 2007; Koivisto et al., 2017).

### Exclusions and Stopping Rule

Participants were excluded from analysis if the total percentage of trials rejected for EEG artifacts (e.g., eye blinks, eye movements, muscle noise, etc.) exceeded 40% for any condition (4 participants were excluded for this reason), or if behavioral responses indicated a failure to perform the experimental tasks as instructed (1 participant was excluded for this reason). Participants who scored lower than 80% on the green circle counting task (i.e., no-report condition) or the animal/object discrimination task (report condition, visible trials only) were also excluded (3 participants were excluded for these reasons). If a participant’s incidental memory test results showed three or more “no” responses for visible stimuli or three or more “yes” responses to either masked or foil stimuli, they were excluded (5 participants were excluded for this reason). Finally, participants who reported seeing the target stimuli on 10% or more of the blank trials in the report condition were excluded (zero participants were excluded for this reason). Data were collected until the total number of non-excluded subjects equaled 20, at which point data collection stopped and data analysis began.

## Results: Experiment 1

### Validation of masking manipulation

“Sandwich masking,” a combination of forward and backward masking, can reliably render stimuli completely invisible or clearly visible depending on the order and timing of the masks and blanks. The current manipulation of stimulus visibility was adapted from Dehaene et al. (2001) and further confirmed in several ways. First, in the report condition, trial-by-trial reports confirmed that the heavily masked stimuli (Figure 2a, right; “masked”) were almost always invisible: participants reported seeing “nothing” on 99.6% (sd=0.5%) of the trials in which animals and objects were masked. The lightly masked stimuli (Figure 2a, left; “visible”) were almost always visible: participants correctly reported seeing “animals” or “objects” on 98.2% (sd=1.8%) of trials. Blank trials were correctly reported as “nothing” on 99.8% (sd=0.3%) of lightly masked trials and 99.7% (sd=0.4%) of heavily masked trials.

**Figure 2.**
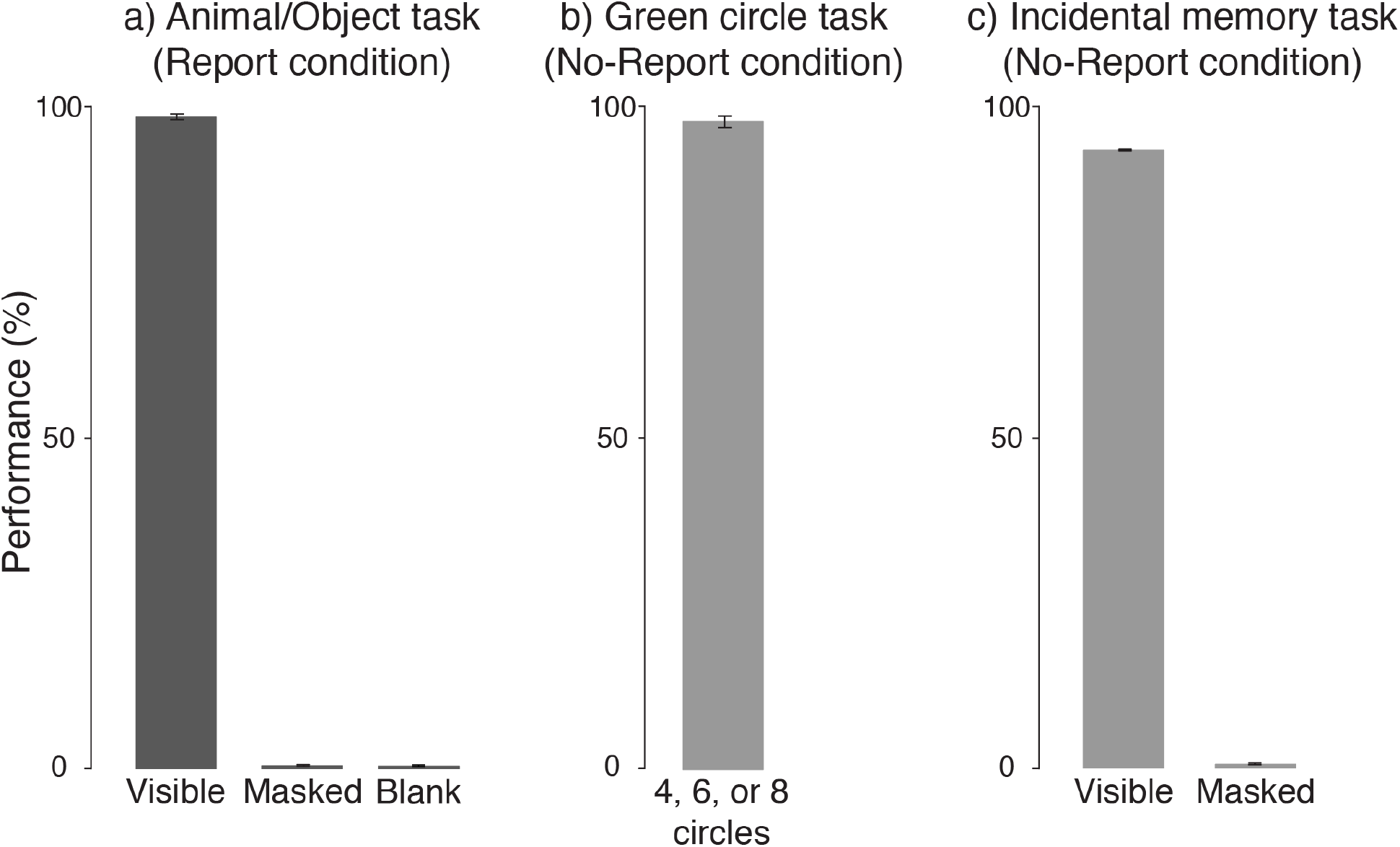
Behavioral results from Experiment 1. In all plots, percent correct (i.e., performance) is plotted on the y-axis). a) Performance on the animal/object/nothing task in the report condition. On the x-axis are the different experimental conditions corresponding to when the target stimulus was visible, invisible (i.e., masked), or absent (i.e., blank). b) Performance on the green circle counting task in the no-report condition. c) Performance on the incidental memory test in the no-report condition for the stimuli that were visible or masked.

Second, on the incidental memory test following the no-report condition, participants responded “yes” 86.3% (sd=10.7%) of the time for stimuli that were visible, 3.8% (sd=8.2%) of the time for stimuli that were masked, and 3.1% (sd=5.5%) of the time for foil stimuli (Figure 2c). Thus, overall accuracy was high (93.1%, sd=4.9%), despite these stimuli being task-irrelevant. After the task-relevant condition, a second memory test was conducted in which subjects reported that they remembered seeing 98.1% (sd=4.6%) of the visible stimuli, 0.6% (sd=2.8%) of the masked stimuli, and 0% (sd=0%) of the foil stimuli. Importantly, across both memory tests, there was no significant difference between “seen” reports for masked stimuli and foil stimuli (*t*(19)=.27, *p*=0.79), further confirming that the masked stimuli were indeed perceptually invisible.

Two important caveats should be made regarding the results from the incidental memory tests. First, these results may somewhat overestimate how frequently participants were aware of the critical stimuli because each individual stimulus was repeated multiple times. In principle, participants could have responded “yes, I remember seeing that stimulus” during the incidental memory test even though they only perceived that stimulus one time out of several presentations. One might reason that this possibility could be mitigated by presenting each critical stimulus only once, while expanding the stimulus set to maintain the same total number of trials. The downside of that approach, however, is that with so many unique stimuli being presented a single time, with no explicit memory task to motivate participants to attend to each individual item, participants’ awareness would likely be artificially underestimated. Even if participants perceived every single stimulus during the experiment, they may be unable to remember many of them and would have difficulty distinguishing them from foil stimuli when later tested. Second, we did not obtain any metacognitive reports from participants regarding their subjective confidence in their responses on the memory test. However, by including foil stimuli in these tests, we were able to assess subjects’ response biases, and observed a very low rate of false alarms. Nevertheless, in the future, acquiring metacognitive data on such memory tests and exploring the trade-offs between a small stimulus set with repeat presentations and a large stimulus set with single presentations could be useful in further characterizing participants’ perceptual experiences in no-report paradigms.

### Hypothesis-driven analyses of the P3b

The main pattern of results was visualized by plotting ERP voltage distribution maps (visible minus masked) from a series of time-windows over all electrode locations and ERP waveforms from a pool of electrodes corresponding to the P3b ROI over all time-points (Figure 3). While early visual ERPs (P1/N1/posterior-P2) were clearly present in both the report and no-report conditions, a sharp divergence was apparent at later time-points, with positive-going ERPs over the fronto-parietal scalp (anterior P2/P3b) being uniquely present in the report condition (grand averaged waveforms from all 64 electrode locations are available on the OSF: https://osf.io/rgfxy/). The observed sequence of ERPs differentiating visible versus masked stimuli in the report condition closely replicated the results of Dehaene et al. (2001). The unique result of the current study was the disappearance of the later ERP effects, including the P3b, in the no-report condition. These robust, late positive ERPs in the report condition vanished in the no-report condition despite the fact that the same stimuli and same manipulation of visual awareness was employed. For the following hypothesis-driven statistical analyses, we quantified the P3b component in each condition by computing the mean voltage from 300-600ms in a pool of 9 electrodes centered around CPz (Pz, P1, P2, CPz, CP1, CP2, Cz, CP3, CP4), a typical spatial-temporal region-of-interest in studies of the P3b (Sergent et al., 2005; Azizian et al., 2006; Del Cul et al., 2007; Polich, 2007; Koivisto et al., 2017).

**Figure 3.**
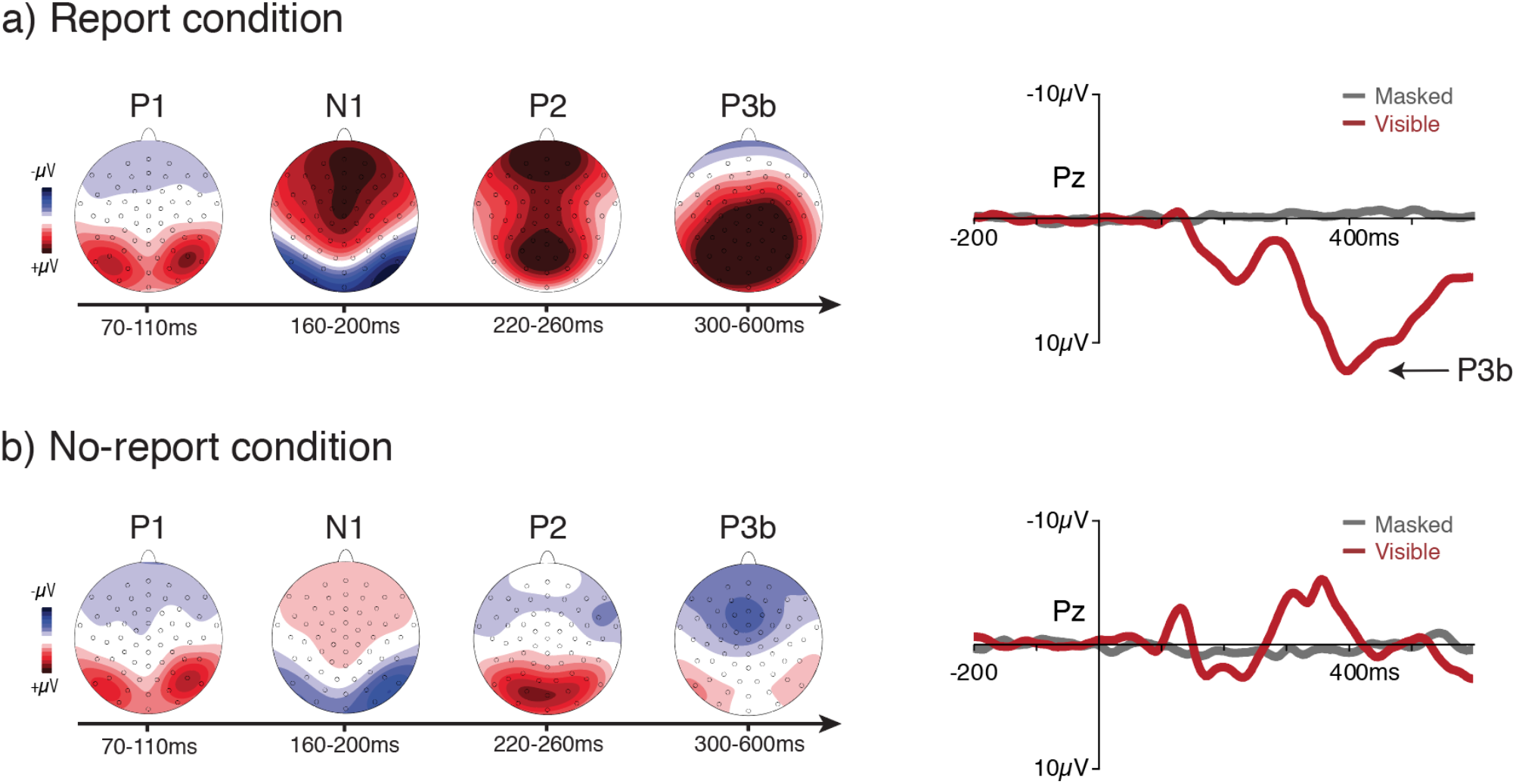
ERP results for both the a) report (top row) and b) no-report (bottom row) conditions. For both condition, topographical voltage distributions over a series of time windows (difference between visible and masked) and the waveforms (for both visible and masked stimuli) from a pool of central-parietal electrodes are plotted. A clear P3b was present in the report condition when observers were aware of the task-relevant stimulus, but the P3b completely vanished in the no-report condition when these same stimuli were task-irrelevant. Amplitude scales for the topography maps: +/- 4μV (P1); +/- 5μV (N1/P2); +/- 6μV (P3b in report condition); +/- 4μV (P3b in no-report condition).

To test the predicted interaction between stimulus visibility and task-relevance, we conducted a 2×2 repeated measures ANOVA with the amplitude measured in the P3b ROI as the dependent variable and with stimulus visibility (i.e., visible versus masked) and reporting task (i.e., report versus no-report) as factors. Main effects of both stimulus visibility (*F*(1,19) = 69.1, *p*<0.001) and reporting task (*F*(1,19) = 95.8, *p*< 0.001) were observed, as was the expected interaction between stimulus visibility and reporting task (*F*(1,19) = 151.13, *p*<0.001). To explore this predicted interaction, we conducted two planned comparison t-tests. In the no-report condition, a one-tailed paired samples t-test showed no significant difference in voltage in the P3b ROI for visible (M = −0.39μV, sd = 1.45μV) versus masked (M = 0.11μV, sd = 0.71μV) stimuli (*t*(19) = −1.51 *p*= 0.93). This same comparison in the report condition confirmed that voltages in the P3b ROI were significantly larger for visible (M = 7.98μV, sd = 3.03μV) as compared to masked stimuli (M = −0.43μV, sd = 1.21μV), (*t*(19) = 11.56, *p*<0.001). Taken together, these results confirm the presence of a P3b in response to stimuli that are visible and reported, but fail to provide evidence for the occurrence of a P3b when these same stimuli are not reported.

Because our hypotheses included the absence of a P3b in the no-report condition, we also conducted planned Bayesian analyses to determine whether our non-significant result was due to cal insensitivity, or whether this result provides evidence in favor of the null hypothesis. First, we compared the mean voltage in the P3b ROI in the no-report condition against a half *t*-distribution with the observed P3b voltage and standard deviation from the report condition as our priors. This analysis returned a BF(10) of 0.001, indicating strong evidence in favor of the null. Next, given that the P3b, if present in the no-report condition, might be substantially smaller than in the report condition, we compared our no-report results to a uniform distribution, specifying 1μv as our minimum effect of interest and our observed voltage for the P3b from the report condition (which was 8 times larger) as the maximum voltage that might be expected. This analysis returned a BF(10) of 0.000, again indicating strong evidence in favor of the null hypothesis, that is, the absence of a P3b for visible yet task-irrelevant stimuli.

### Data-driven analyses

In addition to the planned ANOVA and Bayesian analyses that focused on typical P3b ROIs, we also conducted non-parametric mass univariate analyses (Groppe et al., 2011) across all electrodes and time points to detect any effects outside of this ROI (Figure 4). The false discovery rate (FDR) method was applied to control for multiple-comparisons, ensuring that 5% or less of the significant effects for each mass univariate test are actual false discoveries (Benjamini & Yekutieli, 2001).

**Figure 4.**
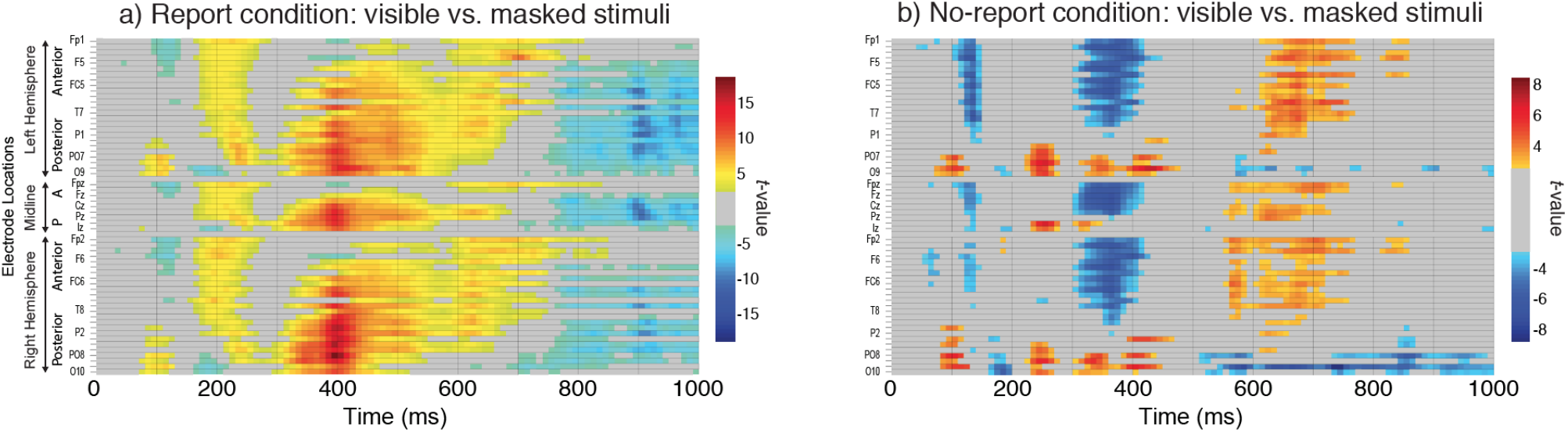
Results from the mass univariate analyses for both (a) the report and (b) the no-report condition. Each individual electrode is plotted as a row on the y-axis, while time (ms) is plotted on the x-axis. Only significant *t*-values (5% FDR) are plotted in the figure.

In the both the report and no-report condition, the contrast between visible and masked stimuli revealed a spatio-temporal sequence of amplitude differences spanning from early (P1: 70-110 ms; anterior N1: 100-140 ms; posterior N1: 160-200 ms; P2: 220-260 ms) to late (fronto-central positivity from 600-750 ms) stages of processing. The P3b was only present in the report condition, evident here as a widespread parietal-central difference between 300-600 ms, maximal at 400 ms. Also unique to the report condition was an earlier fronto-centrally distributed difference from 170-250 ms (anterior P2 or “selection positivity”), likely an attentional effect related to the discrimination task (Hillyard & Anllo-Vento, 1998), and a very late wide-spread negativity (750-1000 ms). Unique to the noreport conditon was a fronto-centrally distributed negative difference from roughly 300-420 ms.

## Materials and Methods: Experiment 2

The main pattern of results from Experiment 1 was robust and unambiguous: the large P3b in the report condition completely disappeared in the no-report condition. However, critics might still question several features of our experimental design and our measures of visibility in the no-report condition. For example, were the “visible” stimuli clearly consciously perceived in the no-report condition (Overgaard & Fazekas, 2016; Phillips, 2018)? After all, these stimuli were task-irrelevant and presented between two masks. Similarly, did our incidental memory test really confirm that subjects consciously perceived the task-irrelevant stimuli in the no-report condition? Perhaps high performance on the memory test could be explained by reactivation of a latent iconic memory trace or by the stimuli in the memory test eliciting a vague sense of familiarity, despite subjects having not consciously seen the stimuli or having only partially seen them (as in seeing oriented lines but not distinct drawings of animals or objects).

To address these concerns, we designed a follow-up experiment. In Experiment 2, the masks were completely removed to enhance visibility of the stimuli in the no-report condition. We also modified the incidental memory test to include words as well as pictures. If participants performed poorly on this conceptual (word) memory test it would be possible to conclude that participants either (a) did not consciously see the stimuli or were only partially aware of the stimuli, or (b) that participants were briefly, phenomenally aware of the stimuli, but failed to access and store their meaning for later conceptual recall. However, if participants performed well on the conceptual memory test, we would be able to safely conclude that the abstract meanings of these stimuli were accessed and stored in episodic memory which is widely believed to be dependent on consciousness. Thus, a combination of good performance on the conceptual memory test and an absence of the P3b in the no-report condition would provide strong evidence against the proposal that the P3b is a marker of conscious awareness.

### Participants

Twenty one participants from the Reed College community completed the experiment in order to obtain a sample size matching Experiment 1. All participants had normal or corrected-to-normal visual acuity, with no head trauma or concussions within the past year and no known history of seizures or epilepsy. Participants were 18-24 years old (mean = 20.2years), 9 male, 12 female, 21 right-handed. Informed consent was obtained prior to each experimental session and all procedures were approved by the IRB at Reed College (IRB no. 2018-S14).

### Stimuli

Experiment 2 used the same stimuli and design as the masking experiment, except that the masks were removed (Figure 5). This necessitated two further changes: (1) Only 8 blocks of trials were presented instead of 15, due to the previously masked stimuli now becoming visible and thus providing twice as many trials per block for analysis (50% stimuli, 50% blanks). Thus, in Experiment 2, the same exact stimuli used in Experiment 1 were presented, but now all 32 stimuli were visible (16 animals, 16 objects), 16 in the noreport condition (8 animals, 8 objects) and 16 in the report condition (8 animals, 8 objects). The same Williams Latin square approach was used to counterbalance stimuli between the report and no-report conditions across subjects. (2) The small red fixation dot, which was present throughout the trials in the masking experiment, was presented here for a variable duration of 200-400 ms before disappearing until the end of the trial. Immediately following the disappearance of the fixation dot, either a stimulus (line drawing of object or animal), or a blank screen, appeared for 33 ms, followed by a blank screen until the end of the trial (total trial duration: 1533-1733 ms). This change allowed the blank screens to signal the viable response window for the discrimination task (3AFC: animal/object/nothing) in the report condition.

**Figure 5.**
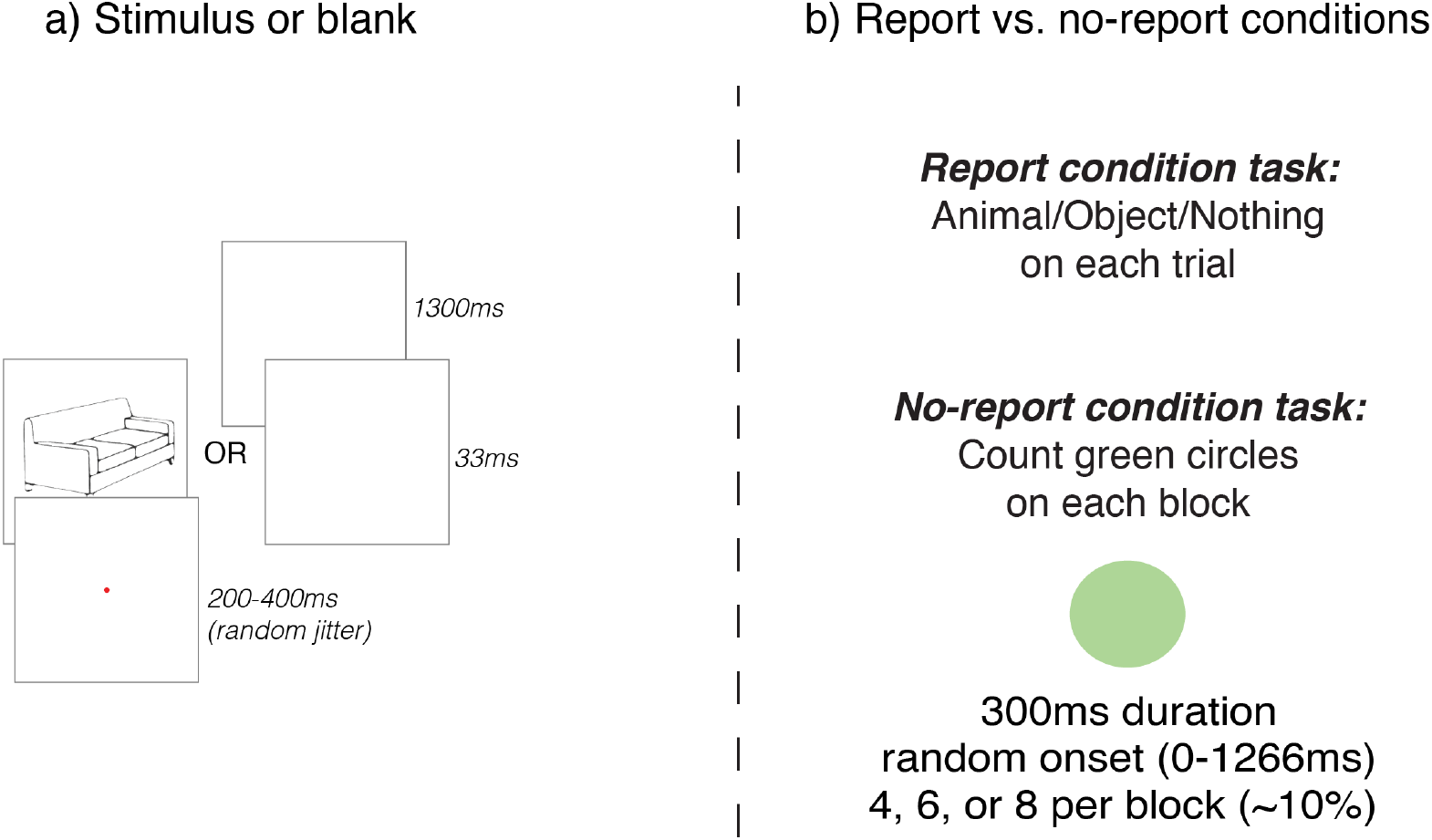
Design of Experiment 2. a) Stimuli (i.e., animals or objects) or blank displays were presented immediately after the fixation dot disappeared. b) In the report condition, participants reported on a trial-by-trial basis whether they saw an animal, object, or nothing. In the no-report condition, the stimulus presentation sequence was the same, but instead of reporting on these stimuli, participants counted the number of times they saw a green circle and reported their count at the end of each block.

### Experimental Design

All procedures were identical to Experiment 1 with the exception of an additional component to the incidental memory tests. Prior to the picture recognition memory test (which was identical to the masking study), participants were presented with words that named the 8 animals and 8 objects that were presented during the experiment, as well as words describing 8 foil stimuli that never appeared. The task was for subjects to report “yes” or “no” as to whether they remember seeing the pictures during the experiment corresponding to each word.

### Statistical Analysis

All recording parameters and ERP analyses were identical to Experiment 1, with the exception that the mask subtraction procedure was no longer necessary, due to the removal of the masks rendering all stimuli visible. On average, 17.8% of trials were excluded due to artifacts in Experiment 2, leaving ~190 trials for analysis per condition. More specifically, in the report condition of Experiment 2, 13.8% of trials were removed for eye blinks and 1.2% for eye movements. In the no-report condition, 15.0% of trials were removed for eye blinks and 1.3% were removed for eye movements. There were no significant differences in the percentage of trials removed for eye blinks or eye movements across conditions (eye blinks: *t*(19)=.639, *p*=0.53, eye movements: *t*(19)=0.59, *p*=0.06). One participant was excluded from analysis due to excessive alpha, resulting in a final sample size of 20.

## Results: Experiment 2

### Incidental Memory Tests

On the conceptual (word) memory test following the no-report condition, participants responded “yes” 69.4% (sd=16.1%) of the time to words naming animals and objects that appeared during the experiment and 3.1% (sd=5.6%) of the time to foil stimuli. Thus, overall accuracy on this memory test was quite high (78.5%, sd=10.9%), suggesting that not only did subjects consciously perceive the task-irrelevant stimuli in the no-report condition, but they encoded many of these stimuli into memory at an abstract conceptual level. On the memory test following the report condition, subjects reported “yes” 86.3% (sd=13.7%) of the time for words naming presented stimuli and 1.9% (sd=4.6%) of the time for foil words (overall accuracy=90.2%, sd=9.9%).

On the picture recognition memory test following the no-report condition, which was always conducted after the word memory test, participants responded “yes” 75.9% (sd=14.7%) of the time for the animals and objects that were presented, and only 0.6% (sd=2.6%) of the time for foil stimuli (overall accuracy = 83.8%, sd=9.8%). In the picture memory test following the report condition, subjects answered “yes” 92.2% (sd=8.8%) of the time for presented stimuli and 1.3% (sd=3.8%) of the time for foils (94.3% accuracy overall, sd=6.1%).

While performance was lower on the word than on the picture memory test, which was expected, we interpret the strong performance on the word memory test as evidence that participants were processing stimuli at a fairly deep level, successfully encoding them into memory, despite them being completely task-irrelevant in the no-report condition. Crucially, participants were highly successful in both memory tests at rejecting foil stimuli, suggesting that participants were not merely guessing or relying on a vague sense of familiarity.

**Figure 6.**
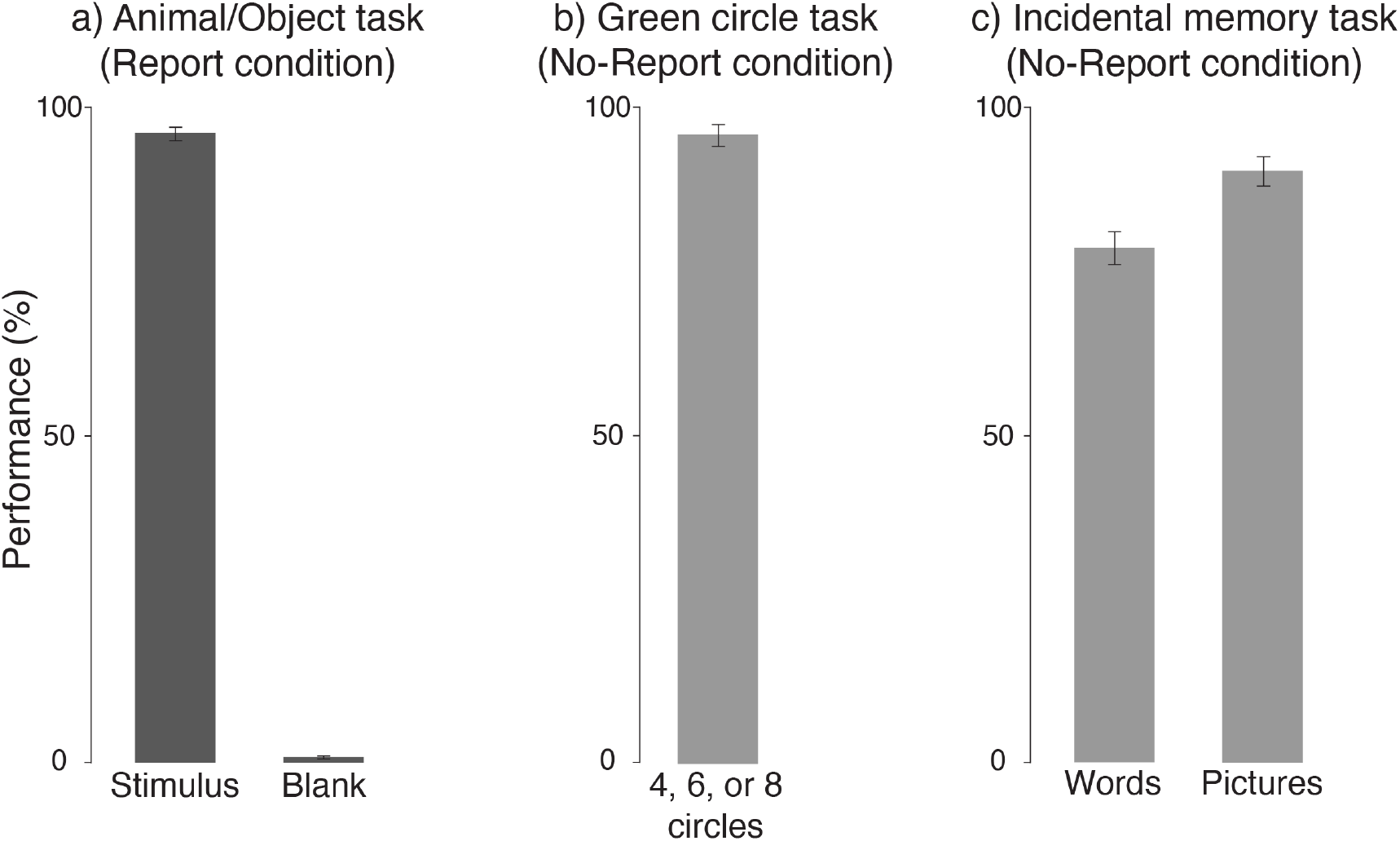
Behavioral results from Experiment 2. In all plots, percent correct (i.e., performance) is plotted on the y-axis. a) Performance on the animal/object/nothing task in the report condition. On the x-axis are the different experimental conditions corresponding to when the target stimulus was present or absent (i.e., blank). b) Performance on the green circle counting task in the no-report condition. c) Performance on the incidental memory task in the no-report condition for both word and picture stimuli.

When comparing the incidental memory test results across the two experiments, it is important to keep in mind that removal of the masks in Experiment 2 resulted in 8 animals and 8 objects being visible in each block (twice as many as in Experiment 1), while each stimulus was presented roughly half as many times, due to the reduction in the number of blocks from 15 (Experiment 1) to 8 (Experiment 2). In other words, participants were tested on twice as many task-irrelevant stimuli, each presented half as many times, which likely accounts for the slight reduction in memory performance in Experiment 2 relative to Experiment 1.

### Hypothesis-driven analyses of the P3b

In Experiment 2, blank trials served as a surrogate for the masked trials in Experiment 1, thus our independent variables were stimulus presence (stimulus, blank) and reporting task (report, no-report). A nearly identical pattern of results to Experiment 1 was observed and is summarized in Figure 7.

**Figure 7.**
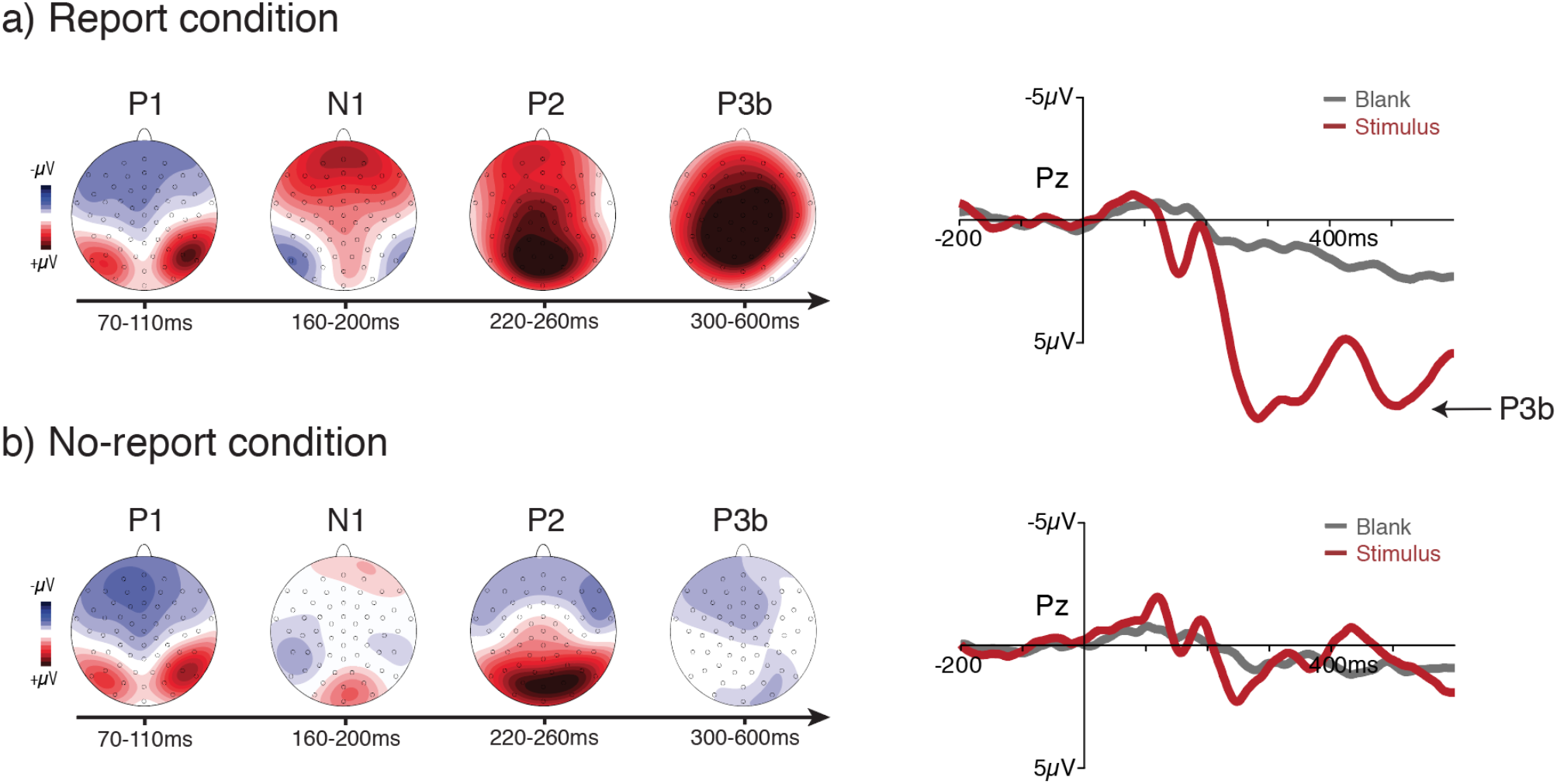
ERP results for both the a) report (top row) and b) no-report (bottom row) conditions. For both conditions, topographical voltage distributions over a series of time windows (difference between visible and masked) and the waveforms (for both visible and masked stimuli) from a representative electrode (Pz) are plotted. A clear P3b was present in the report condition when observers were presented a task-relevant stimulus, but the P3b completely vanished in the no-report condition when these same stimuli were task-irrelevant. Amplitude scales for the topography maps: +/- 4μV (P1); +/- 5μV (N1/P2); +/- 4μV (P3b). Note that in this experiment, a posterior N1 was elicited by both stimuli and blanks (fixation dot offset), thus N1 amplitude differences (stimulus minus blank) were much smaller than in Experiment 1.

Significant main effects of stimulus presence (F(1,19) = 37.0, *p*<0.001) and reporting task (F(1,19) = 49.6, *p*<0.001) were observed, as was the expected interaction between stimulus presence and reporting task (F(1,19) = 58.6, *p*<0.001). Planned-comparisons t-tests confirmed that this interaction was driven by larger voltages in the P3b ROI for stimuli (M = 6.55μV, sd = 3.62μV) versus blanks (M = 1.78μV, sd = 1.80μV) in the report condition (t(19) = 8.30 *p*<0.001), and that there was no significant difference in voltages for stimuli (M = 0.462μV, sd = 1.76μV) versus blanks (M = 0.748μV, sd = 1.20μV) in the no-report condition (t(19) = −0.71 p = .758).

The same Bayes factor analyses used in Experiment 1 indicated evidence in favor of the null hypothesis: A comparison of P3b voltage in the no-report condition versus a half t-distribution, specified using the P3b voltage and standard deviation in the report condition, yielded a BF(10) of 0.05. Similarly, comparison of P3b voltage in the no-report condition versus a uniform distribution with 1μv as the minimum effect of interest and our observed voltage for the P3b in the report condition (6.5μV) as the maximum voltage that might be expected, returned a BF(10) of 0.03, again indicating evidence in favor of the null hypothesis, that is an absence of a P3b in the no-report condition.

### Data-driven analyses

Similar to Experiment 1, in both the report and no-report conditions, amplitude differences were evident during early (P1; anterior N1; posterior N1; P2) and late (fronto-central positivity and posterior negativity, 600-800 ms) time windows. Unique to the report condition, the P3b was evident as a widespread positive difference from 300-600 ms, and we again observed an earlier fronto-centrally distributed difference from 150-200 ms for stimuli versus blanks, likely related to attentional demands of the discrimination task. Unique to the no-report condition, the contrast between stimuli and blanks revealed a fronto-central negativity from 420-500 ms. These results are shown in Figure 8.

**Figure 8.**
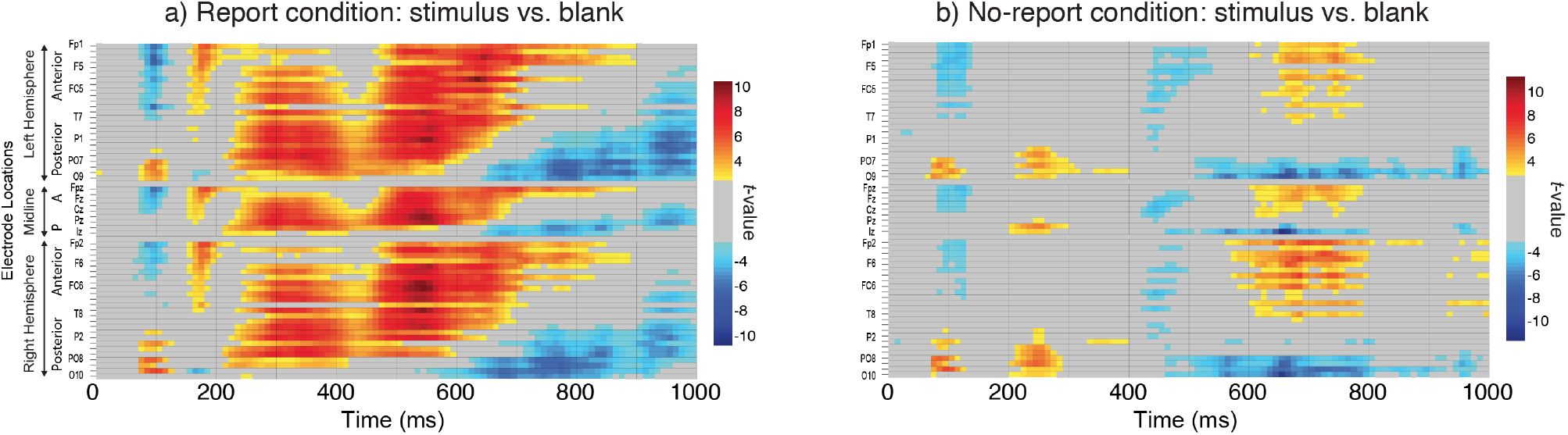
Results from the mass univariate analyses for both (a) the report and (b) the no-report condition. Each individual electrode is plotted as a row on the y-axis, while time (ms) is plotted on the x-axis. Only significant *t*-values (5% FDR) are plotted in the figure. Note that the absence of a posterior N1 (160-200ms) was due to both stimulus and blank trials eliciting an N1.

### Results summary

The pattern of results from Experiment 2 closely replicated the results from Experiment 1, while alleviating several remaining concerns about stimulus visibility in the no-report condition. Indeed, the P3b remained completely absent in the no-report condition, even without masks that could potentially disrupt perception, and despite strong performance on the word-based (conceptual) memory test suggesting that the task-irrelevant stimuli in the no-report condition were consciously accessed and encoded in an abstract format.

## What happens after or instead of the P3b?: Exploratory analyses

The focus of these experiments was to examine the interaction between stimulus visibility and task relevance on the P3b. After finding clear evidence that the P3b is only present when stimuli are relevant to the task, we then examined our data driven results (Figures 4 and 8) to explore any commonalities across the report and no-report conditions as well as other effects that were unique to one condition or the other.

As mentioned above, in both the report and no-report conditions, visible versus masked stimuli elicited differential ERP amplitudes during early time windows from ~70-260 ms (P1, N1, P2). This is not surprising given the heavy masking used in this experiment, which most likely blocked early stages of sensory processing for masked relative to visible stimuli. Unexpected, however, was the presence of amplitude differences in both the report and no-report conditions during late time windows, from ~600-800 ms (Figure 9). These differences were characterized by a fronto-central positivity coupled with a bilateral posterior negativity. Even later in time (800-1000 ms), posterior negativities were observed in both conditions. The scalp distributions of these late-stage effects were clearly distinct from the P3b and should be investigated further in future studies as they persisted regardless of the reporting task.

**Figure 9.**
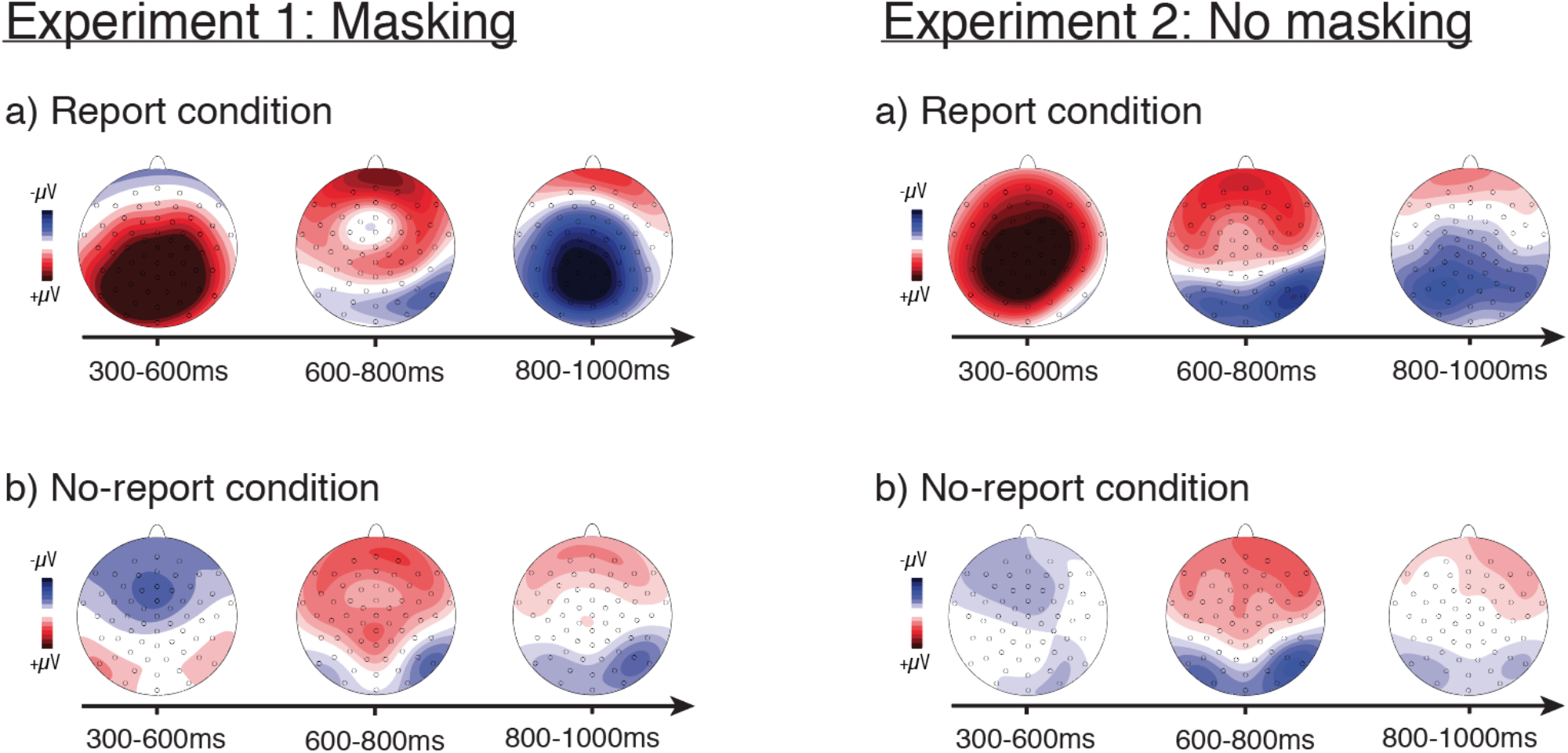
ERP differences during late time windows revealed by exploratory analyses for both Experiment 1 (visible versus masked stimuli; left panels) and Experiment 2 (stimuli versus blanks; right panels). For both experiments, topographical differential voltage distributions over a series of time windows starting at 300ms and ending at 1000ms are shown for both report (top) and no-report (bottom) conditions. Amplitude scales for the topography maps: +/- 6μV (300-600ms in the masking report condition); +/- 4μV (all other maps).

Another unexpected result was a fronto-central negativity from ~300-500 ms that was evident only in the no-report condition (Figure 9). Indeed, the data driven statistics (Figure 4 and 8) suggest that this ERP difference occurred within a similar time frame as the P3b in the report condition and could reflect a task-related effect unique to the no-report condition. Possible functional interpretations of this fronto-central negativity should be investigated in future experiments. Taken as a whole, these data-driven exploratory analyses appear to indicate strong task-driven ERP effects during the 300-600 ms timewindow, with similarities across report and no-report conditions both earlier and later in time.

## Source estimations during late time windows: Exploratory analyses

In order to further explore the results depicted in Figure 9, we next estimated the anatomical sources of the differential brain activity in the visible vs. masked conditions from Experiment 1. All source estimations were performed using Brainstorm (Tadel et al. 2011). Here, our electrode coordinates were mapped to the anatomy of the MNI/ICBM52 brain and a forward model was generated via OpenMEEG BEM (Gramfort et al. 2010, Kybic et al. 2005). Source current density maps were then estimated via minimum norm imaging using an unconstrained source model. Average activation from 300-600 ms, 600-800 ms, and 800-1000 ms in each condition was parametrically tested against zero with control of the false discovery rate set at the *p*<0.05 significance level. The results from these analyses are plotted in Figure 10.

**Figure 10.**
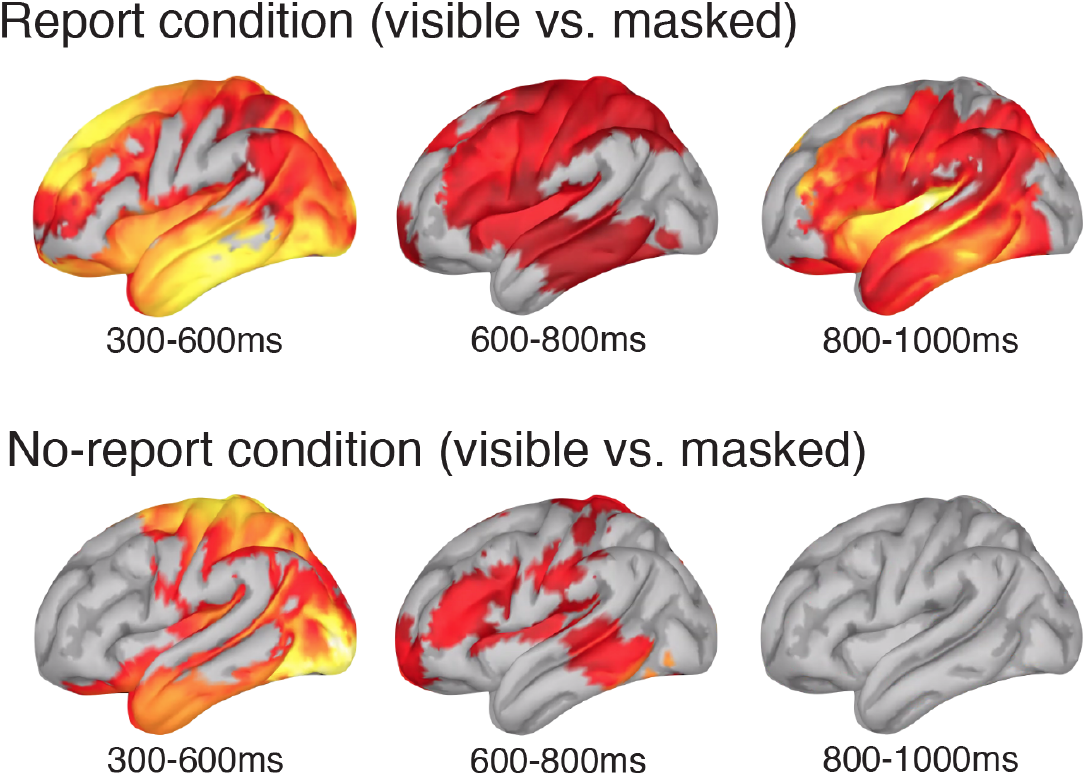
Source estimations of the difference between visible and masked stimuli in Experiment 1 for both the report (top row) and no-report (bottom row) conditions across three late time windows (see Figure 9 for the average voltage distributions during these same time windows). Only significant *t*-values (5% FDR) are plotted on the lateral surface of the Montreal Neurological Institute brain.

It is worth noting that these source estimates were entirely exploratory and were not pre-registered on the Open Science Framework. In addition, the experiments reported here were specifically designed to examine the role of the P3b in perceptual awareness, not to identify or isolate other potential neural correlates of perceptual awareness. With these qualifications in mind, these exploratory analyses suggest a few noteworthy trends that could potentially be explored in future research. First, between 300-600 ms, during the time period of “ignition” predicted by the global neuronal workspace theory (Mashour et al., 2020), there was a clear decrease in estimated source activity in the frontal lobe in the noreport condition relative to the report condition. However, between 600-800 ms, there was a small amount of estimated source activity in the frontal lobe in the no-report condition, though it was still reduced compared to the report condition. Second, although there was little to no estimated activity in the frontal lobe between 300-600 ms in the no-report condition, there were several areas within the parietal lobe showing estimated sources during this time window. While the frontal and parietal lobes are often treated as a singular entity in consciousness studies (i.e., “the fronto-parietal network”), these findings tentatively suggest that these two regions may have dissociable roles in perceptual awareness. Finally, as far out as the 800-1000 ms time window, there is considerably more estimated activity in the report condition than in the no-report condition. Once again, it should be stressed that each of these results should be interpreted with extraordinary caution. Source estimations from ERP data are notoriously imprecise, particularly in experiments such as ours in which the experimental design was not optimized to isolate sources (e.g. the contrast between visible and masked stimuli employed here likely overestimates neural differences between conscious and unconscious processing). Going forward, it will be important to explore these issues with methods like functional magnetic resonance imaging (fMRI) and with paradigms that better isolate the perceptual content of which subjects are aware versus unaware.

## Discussion

Here, we found strong evidence suggesting that the P3b is not a signature of perceptual awareness per se and is instead more closely associated with post-perceptual processing. While we replicated prior results showing a robust P3b in visible conditions when observers reported their experience (Dehaene et al., 2001), this effect completely disappeared when observers perceived the same stimuli but did not have to report on their experiences (see also Koivisto & Revonsuo, 2008; Pitts et al., 2012; 2014). Furthermore, the P3b appearing and disappearing as a function of task demands was seen both in Experiment 1, where we manipulated visibility with a standard masking procedure, as well as in Experiment 2, where the masks were removed and the stimuli were always visible. Together, these results demonstrate that the P3b is not a true neural correlate of conscious processing and raise a variety of important questions.

Perhaps the most important question is whether we believe that our results disprove or invalidate cognitive theories of conscious awareness? No. It is worth stressing that we are in no way claiming that our data suggests that cognitive and higher-order theories are incorrect and that sensory theories are correct. Cognitive and higher-order theories of awareness may ultimately be correct even if the P3b is not a neural signature of consciousness. This could be the case for several reasons. For example, it is possible that there is significant fronto-parietal activation when participants are aware of the stimuli in these no-report situations but we are simply unable to measure this activation using EEG and the P3b. Given the limits of the spatial resolution of EEG, it could easily be the case that such fronto-parietal activation can only be observed with other methods such as functional magnetic resonance imaging or electrocorticography. In addition, it is also possible that while the P3b is not a signature of conscious processing, some of the other putative signatures (i.e., an “ignition” of fronto-parietal circuits, an increase in long-range neural synchrony, and a late burst of gamma-band oscillations; Dehaene, 2014) could still be found using a no-report paradigm such as ours. Such results would lend strong support for cognitive and higher-order theories even if the proposed P3b signature must be discarded. Thus, we believe future research is needed before any definitive claims can be put forth regarding the long-standing debates on the neural correlates of awareness and between these two classes of theories. Specifically, we maintain that no-report paradigms such as ours should be replicated with other methodologies in order to carefully test for the presence of the remaining neural signatures of perceptual awareness proposed by cognitive theories.

While we firmly believe our results do not critically challenge the core hypotheses of cognitive and higher-order theories, they do challenge certain long-standing, key predictions, particularly of global neuronal workspace theory, which states that the P3b is a signature of conscious processing (Dehaene, 2014). However, before our conclusion that the P3b is *not* a signature of conscious vision can be accepted, there are important questions that must be considered. For example, one important question asks if we can be sure that our observers were aware of the critical stimuli in the no-report condition when we have no objective way to probe their experience on a trial-by-trial basis (Overgaard & Fazekas, 2016; Phillips, 2018). Without such objective measures, how can we be sure that observers perceived the irrelevant stimuli at all?

We believe observers were aware of the critical stimuli in the no-report condition for three reasons. First, we modified the visual masking paradigm such that the amount of time between the critical stimuli and masks in the visible condition was longer than in prior studies in order to maximize the visibility of the stimuli in the visible condition (200 ms in our study versus 71 ms in Dehaene et al., 2001). These efforts appear to have succeeded by virtue of the fact that when reporting the contents of their experience on a trial-by-trial basis, observers were effectively always able to correctly identify the targets (see Figure 2). Second, we conducted an incidental memory test in which observers were unexpectedly asked to retrospectively identify the items they were shown in the no-report blocks specifically to obtain objective measures of their perceptual experience. In these cases, the overwhelming majority of observers were able to correctly identify which items they had and had not been shown in the no-report condition. We believe that observers’ ability to perform this surprise memory task so well was possible because they were aware of the stimuli in the no-report condition. Third, a major motivation for Experiment 2 was to present the stimuli in a manner that would limit the possibility that observers would fail to perceive the stimuli in the no-report condition. As a reminder, Experiment 2 followed the same procedures as Experiment 1 with the only major difference being that we removed the masks. Thus, participants were simply presented with isolated objects with nothing being shown before or after them. In essence, we took all possible steps we could think of to ensure observers would be aware of the stimuli even if we were never directly asking for a report on each trial.

Another question to consider with no-report paradigms is whether our observers engaged in spontaneous post-perceptual processing even in no-report conditions (Block, 2019). Since the task demands of this no-report paradigm were relatively minor (i.e., the green circle counting task required minimal attention and effort), perhaps our observers naturally engaged in a variety of post-perceptual processes even if they never had to report their perceptual experiences. For example, when shown a picture of a penguin, an observer may spontaneously think, “Oh look, a penguin.” And then a few trials later, when shown the same penguin, the observer may think, “That’s the same penguin I just saw. I wonder if I’ll see again.” In that situation, even though we would have eliminated the need for decision-making and motor planning related to that particular trial, we would not have eliminated all post-perceptual processing related to the stimulus. However, for the purposes of this study, we believe it is highly unlikely that such factors affected our findings. If observers naturally engaged in post-perceptual processing in no-report conditions and we still observed a P3b in the no-report condition, this could lead to difficulties in interpretation. If a P3b was observed in no-report conditions, it might index neural activity related to conscious perception or it might reflect neural activity associated with spontaneous cognitive processing. In our data, however, we find no such P3b associated with the critical stimuli in the no-report condition. Thus, we conclude that the P3b is more tightly linked with carrying-out reporting tasks than with consciously perceiving the stimuli. While we do not believe the concern of spontaneous cognitive processing is a problem in our experiments, we certainly agree that it is a difficult and important issue that must be consistently wrestled with when using no-report paradigms (Block, 2019).

After ruling out the P3b, were there any remaining candidate neural signatures of awareness in both the report and no-report conditions? Importantly, the current study was not designed to isolate any such signatures, but was instead aimed at testing specific predictions regarding the P3b. The masking procedure was so severe that early (pre-conscious) sensory activity was disrupted in the invisible condition leading to an overestimation of neural differences between seen and unseen stimuli. Indeed, P1/N1 were larger in visible versus invisible conditions (Figures 3 & 7), but it is well established that P1 does not track perceptual awareness (Railo et al., 2011). The N1 difference might, however, correspond to the visual awareness negativity (VAN) (Ojanen et al., 2003; Koivisto & Revonsuo, 2010; Forster et al., 2020). Later differences, beyond the P3b time window (600-1000 ms), were observed in both the report and no-report conditions as well (Figures 4, 8, 9, & 10). Of particular interest was a fronto-central positivity and posterior negativity from ~600-800 ms that was evident regardless of the reporting task in our exploratory data-driven analyses. Taken together, consistent neural differences between visible and invisible stimuli were evident during both early (70-300 ms) and late (600-1000 ms) time windows, while neural differences between 300-600 ms were highly dependent upon the reporting task.

## Conclusion

Overall, these results challenge the idea that the P3b is a neural signature of conscious processing. While this particular neural activation pattern has been widely replicated and continues to be systematically put forth as a possible neural signature of consciousness (Railo et al., 2010; Dehaene, 2014; Naccache, 2018; Ye et al., 2019; Derda et al., 2019), these prior studies always had participants report their experiences. Although it may be the case that the theoretical foundation supporting those prior studies may ultimately be proven correct (i.e., cognitive/higher-order theories), these results conclusively demonstrate that the P3b is not a signature of conscious processing. Together, these findings highlight the need to differentiate between perceptual awareness and post-perceptual processing in future studies that use different methodologies or analysis methods. No-report paradigms, such as the one employed here, should be considered useful tools for probing the neural underpinnings of conscious perception and for testing various predictions made by scientific theories of consciousness.

## Acknowledgments

We thank Steve Hillyard for his contributions to helping design the no-report paradigms used here, and Sid Kouider for suggesting the conceptual memory test used in Experiment 2. This work was supported by NSF Grant BCS-1829470 to M.A.C. & M.P.

## Notes

### Competing Interest Statement

The authors have declared no competing interest.

